# *Azotobacter vinelandii* scaffold protein NifU transfers iron to NifQ as part of the iron-molybdenum cofactor biosynthesis pathway for nitrogenase

**DOI:** 10.1101/2021.12.06.471366

**Authors:** Emma Barahona, Xi Jiang, Emilio Jiménez-Vicente, Luis M. Rubio, Manuel González-Guerrero

## Abstract

*Azotobacter vinelandii* molybdenum-dependent nitrogenase obtains molybdenum from NifQ, a monomeric iron-sulfur molybdoprotein. This protein requires of a preexisting [Fe-S] cluster to form a [MoFe_3_S_4_] group to serve as specific donor during nitrogenase cofactor biosynthesis. Here, we show biochemical evidence for NifU being the donor of the [Fe-S] cluster. Protein-protein interaction studies using apo-NifQ and as-isolated NifU demonstrated the interaction between both proteins which is only effective when NifQ is unoccupied by its [Fe-S] cluster. The apo-NifQ iron content increased after the incubation with as-isolated NifU, reaching similar levels to holo-NifQ after the interaction between apo-NifQ and NifU with reconstituted transient [Fe_4_-S_4_] groups. These results also indicate the necessity of co-expressing NifU together with NifQ in the pathway to provide molybdenum for the biosynthesis of nitrogenase in engineered nitrogen-fixing plants.

## Introduction

Nitrogenases catalyses the reduction of N_2_ into NH_3_ in a energetically expensive process (1). These enzymes, only present in some bacteria and archaea, are two-component oligomeric metalloprotein complexes made up of a dinitrogenase (component I) and a dinitrogenase reductase (component II) (2). Component I of molybdo-nitrogenases, the most common ones, is a heterotetramer formed by two NifD, two NifK proteins and two different metalloclusters. The iron-molybdenum cofactor (FeMo-co; [Fe_7_-S_9_-C-Mo-*R*-homocitrate]) is present at the active site of each NifD subunit, while the [Fe_8_–S_7_] P-cluster is at the interface of each NifD and NifK subunits (3, 4). Component II is a homodimer encoded by *nifH*. This protein contains a single [Fe_4_-S_4_] cluster bridging the two identical subunits and two sites for Mg^2+^-ATP binding and hydrolysis (1). Electrons provided to NifH are transferred from its [Fe_4_-S_4_] cluster through the P-cluster of NifDK to FeMo-co, where N_2_ is reduced (5, 6). Therefore, for nitrogenase to function, these metal cofactors must be assembled, protected from oxygen, and transferred to the apo-enzymes, a tightly-regulated process that requires several additional proteins (5). Among them, NifU and NifQ are the known points from where iron and molybdenum are specifically directed towards nitrogenase cofactor assembly (5).

NifU is a 33 kDa homodimer with a permanent [Fe_2_-S_2_] cluster per subunit (7). It is able to bind iron to synthesize [Fe_4_-S_4_] groups, using the sulfur provided by NifS, a 43 kDa cysteine desulfurase (8, 9). These groups are transiently assembled in N- and C-terminal domains, and are subsequently transferred to apo-NifH, activating it (10). NifU is also involved in FeMo-co biosynthesis, providing the substrate [Fe_4_-S_4_] clusters required for NifB-co assembly by NifB (11). These data indicate a pivotal role of NifU in [Fe-S] assembly and transfer to the different enzymes involved in nitrogenase maturation and cofactor assembly.

Molybdenum destined for FeMo-co assembly is typically provided by NifQ. *Azotobacter vinelandii* and *Klebsiella pneumoniae nifQ* mutant strains are impaired in nitrogen fixation unless molybdate levels are dramatically increased in the growth medium (12, 13). NifQ is a 22 kDa monomeric [Fe-S] molybdoprotein that may contain three to four iron and up to one molybdenum atoms per molecule (14). This protein is found in all diazotrophic species of Proteobacteria (excepting some *Rhizobia*) (15). Although the mechanism is yet-unknown, it has been shown that NifQ synthesizes a [Mo-Fe_3_-S_4_]^3+^ group using a [Fe_3_-S_4_]^+^ precursor (16). Subsequently, this Mo-Fe-cluster will be transferred to a NifEN/NifH complex for molybdenum integration into FeMo-co (14). Currently, the source of the [Fe-S] cluster precursor of NifQ is unknown.

Considering the central position of NifU as the scaffold in which [Fe-S] clusters are first assembled for some nitrogenase components (10, 11), it can be hypothesized that it is also the source of the NifQ clusters. Supporting this role, here we report that NifU transfers a [Fe_4_-S_4_] cluster to NifQ through direct protein-protein interaction.

## Results

### NifU and NifS co-elute with NifQ

To determine whether NifQ can interact with the [Fe-S] cluster biosynthesis branch of the nitrogenase assembly pathway, an N-terminal Strep-tagged *A. vinelandii* NifQ (_S_NifQ) was expressed in an *Escherichia coli* strain that already produced *A. vinelandii* NifU and NifS proteins. After induction and cell lysis, _S_NifQ was purified under anaerobic conditions. As expected, _S_NifQ was the most abundant protein in the eluted fractions of the StrepTactin Affinity Chromatography (STAC) chromatography, as evidenced by the Coomassie blue staining of SDS-gels as well as the immunodetection of NifQ with specific antibodies (Fig. 1). To determine whether NifU and NifS were among these additional bands, specific antibodies raised against either protein were used for immunoblotting. As shown in Figure 1, both proteins co-eluted with NifQ. These were not the result of unspecific interaction of NifU and/or NifS with the purification resin, since both proteins were not detected in the elution fractions when NifQ was not expressed in this *E. coli* strain (Fig. S1).

**Figure 1.**
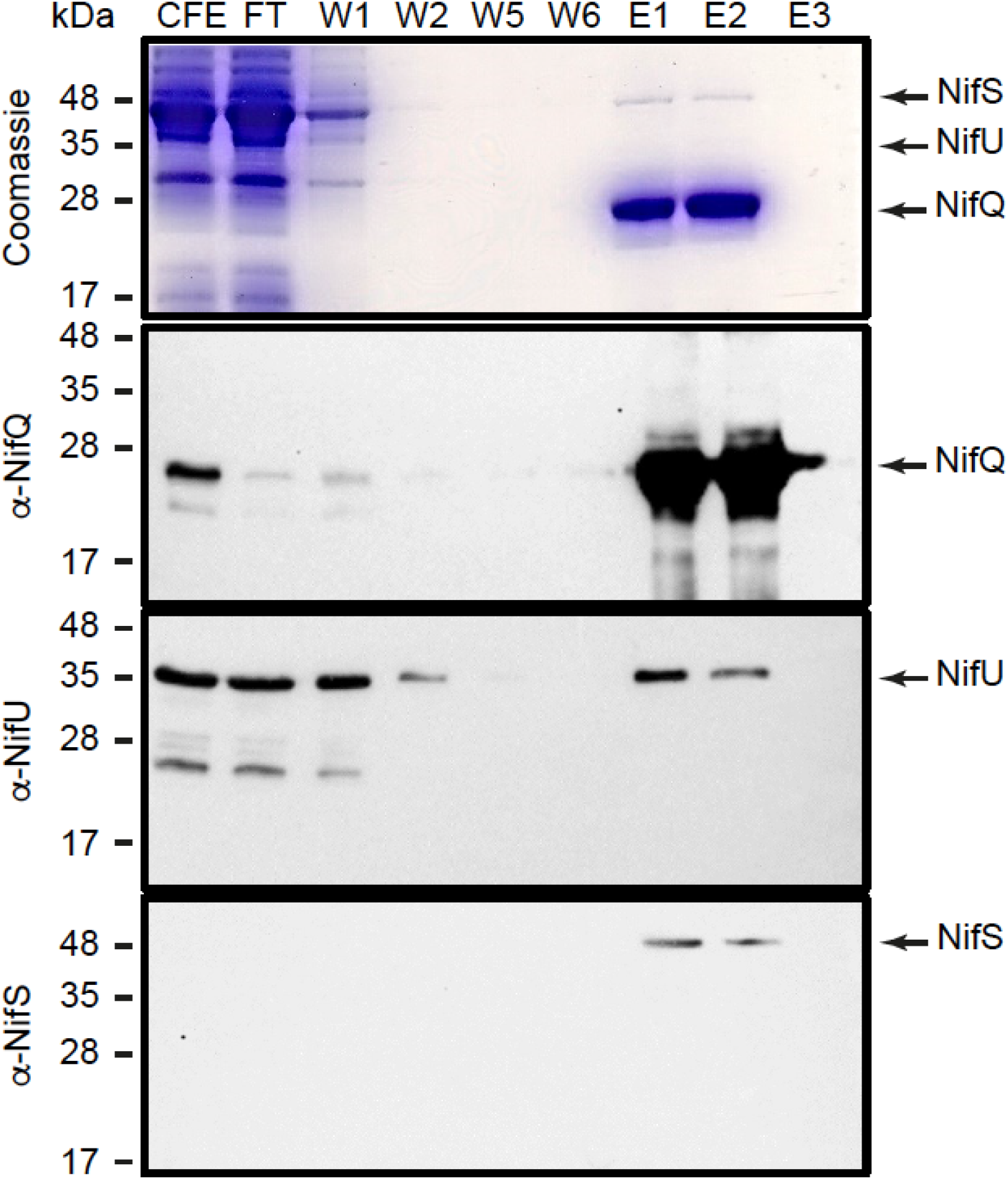
NifU and NifS copurify with _S_NifQ from *E. coli* extracts expressing *nifQ, nifU*, and *nifS*. Top panel shows the Coomassie staining of an SDS-PAGE of cell free extract (CFE), flowthrough (FT), wash (W1-W6) and elution (E1-E3) fractions of the extracts passed through a Streptactin column. The remaining panels show immunoblots of the same fractions developed with anti-NifQ, anti-NifU, or anti-NifS antibodies. Images show a representative assay (n=2). Uncropped immunoblots and gels are shown in Supplementary Fig. 4.

### The interaction between NifQ and NifU is NifS independent and apo-NifQ dependent

The co-purification of NifU and NifS with NifQ from *E. coli* crude extracts could be the consequence of direct interactions among these three proteins, or in combination with endogenous proteins. To discriminate between these two possibilities, apo-NifQ was purified using a (His)_6_ tag (apo-NifQ_H_). This apo-form had less than 1 iron atom per monomer (Table 1). The “*as purified*” (AS) NifU_S_ contained 2.4 iron atoms per monomer (Table 1), a mix of the permanent [Fe_2_-S_2_] cluster and the transient ones. Apo-NifQ_H_, AS-NifU_S_ and _S_NifS were incubated together for 5 minutes under anaerobic conditions and loaded onto Ni-NTA column. As shown in Figure 2, apo-NifQ_H_ was properly captured by the resin and the protein eluted at 150 mM imidazole. Most soluble AS-NifU_S_ was detected in the flowthrough and early wash fractions, but a significant amount co-eluted with apo-NifQ_H_. AS-NifU_S_ presence in the elution fractions was due to apo-NifQ_H_; when NifQ was not present, no NifU was detected in the eluates (Fig. S2). These results confirm the apo-NifQ/AS-NifU interaction without additional proteins being required. On the contrary, _S_NifS was only detected in the flowthrough and initial wash fractions (Fig. 2), suggesting that NifS was not necessary for the apo-NifQ/AS-NifU interaction.

**Table 1.** Proteins used in this work. Data are the average iron content per protein monomer ± SD calculated for apo-_S_NifQ (n=3), apo-NifQ_H_ (n=8), holo-_S_NifQ (n=2), AS-_H_NifU (n=2), AS-NifU_S_ (n=2), and R-NifU_S_ (n=2). S indicates Strep-tagged protein; H, 6xHis-tagged; AS, as isolated protein; and R, reconstituted [Fe-S] clusters.

| Protein | Name | Tag/ Position | Fe/ monomer | Source |
| --- | --- | --- | --- | --- |
| NifQ | apo-sNifQ | Strep/ N-t | $0.56 \pm 0.01$ | This work |
| | apo-NifQ<sub>H</sub> | 6xHis/ C-t | $0.04 \pm 0.01$ | This work |
| | holo-sNifQ | Strep/ N-t | $2.82 \pm 0.19$ | This work |
| NifU | AS-<sub>H</sub>NifU | 6xHis/ N-t | $2.72 \pm 0.46$ | (28) |
| | AS-NifU<sub>S</sub> | Strep/ C-t | $2.38 \pm 0.28$ | This work |
| | R-NifU<sub>S</sub> | Strep/ C-t | $5.82 \pm 0.62$ | This work |
| NifS | sNifS | Strep/ N-t | - | This work |

**Figure 2.**
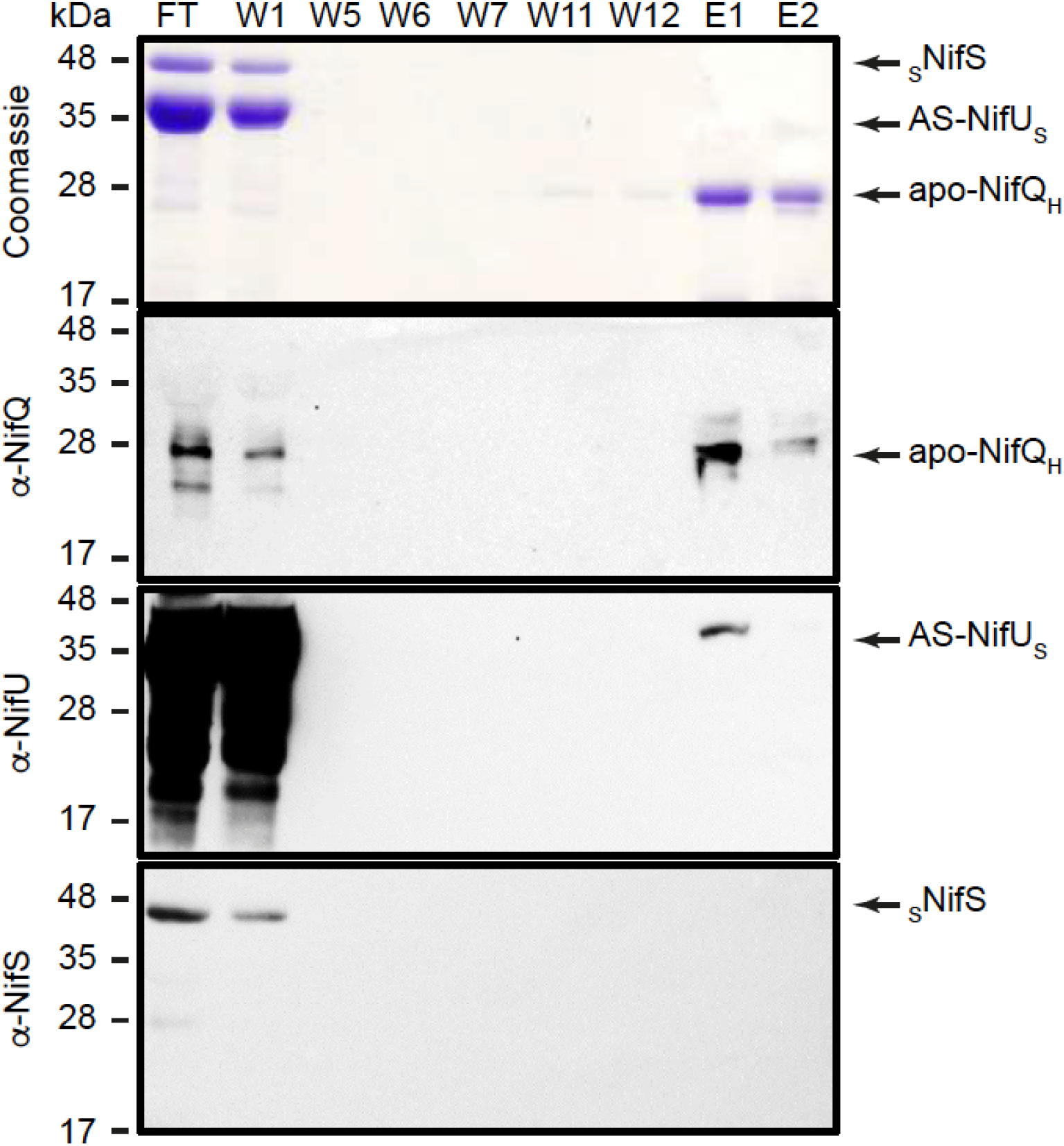
AS-NifU_S_ interacts with apo-NifQ_H_. Top panel shows the Coomassie staining of an SDS-PAGE of flowthrough (FT), wash (W1-W12) and elution (E1-E2) fractions of a mixture solution containing AS-NifU_S_, _S_NifS, and apo-NifQ_H_ passed through a Ni^2+^ column. The remaining panels are the immunoblots of the same fractions developed with anti-NifQ, anti-NifU, or anti-NifS antibodies. Images show a representative assay (n=3). Uncropped immunoblots and gels are shown in Supplementary Fig. 5.

Considering that the metalation state of NifQ might influence the interaction with NifU, co-purification assays were carried out between _S_NifQ in its holo-state and a N-terminal (His)_6_-tagged NifU (AS-_H_NifU). Holo-_S_NifQ had 2.8 iron atoms per monomer and AS-_H_NifU had 2.7 (Table 1). In contrast to what was observed using apo-_S_NifQ, no interaction with NifU was observed (Fig. 3). This data suggests that when NifQ is already occupied by an [Fe-S] cluster the interaction with NifU is much reduced.

**Figure 3.**
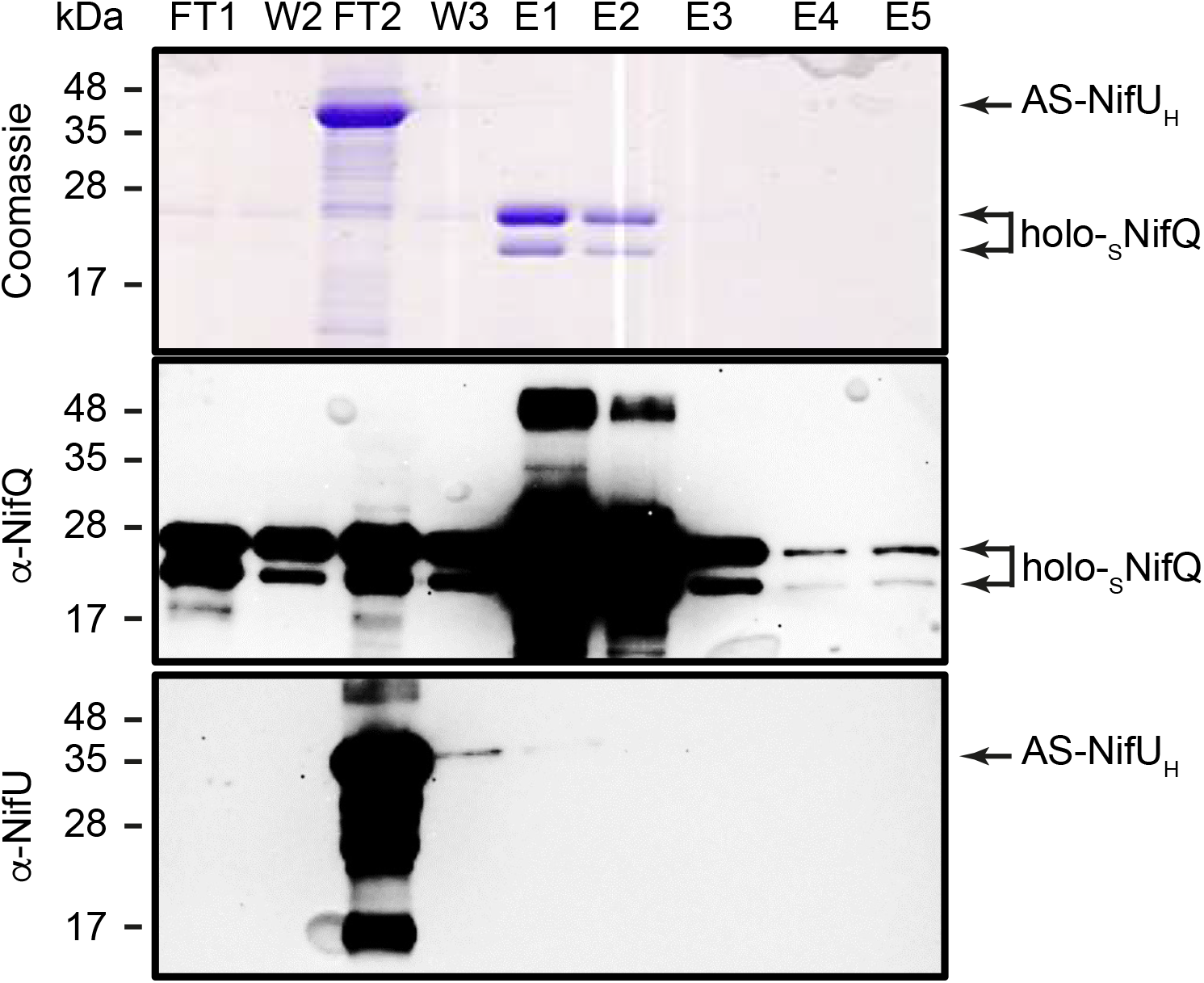
AS-_H_NifU does not co-elute with holo-_S_NifQ. Top panel shows the Coomassie staining of an SDS-PAGE of AS-_H_NifU pull-down assay using holo-_S_NifQ as bait. FT1 represents flow-through fraction obtained after loading holo-_S_NifQ onto the column. W2 represents the second wash fraction. FT2 represents flow-through fraction obtained after loading AS-_H_NifU onto the column. W3 represents the third wash fraction after passing AS-_H_NifU over holo-_S_NifQ-charged column. E1, 2, 3, 4 and 5 represent elution fractions. (*n*=4). Uncropped immunoblots and gels are shown in Supplementary Fig. 6.

### The iron content of NifQ increases after the interaction with NifU

The fact that the interaction between NifQ and NifU is contingent upon the iron content of NifQ is indicative of a process in which iron would be transferred from NifU. This possibility was tested by determining the iron transfer from one protein to the other. Apo-NifQ_H_ was incubated with either AS-NifU_S_ or with a reconstituted (R) NifU_S_ that contained a higher complement of transient [Fe_4_S_4_] clusters, as indicated by a 5.8 iron:monomer ratio (Table 1). These proteins were incubated for 5 min to allow for metal transfer. The interaction with R-NifU_S_ did not seem to be more stable than with AS-NifU_S_, since similar amounts of proteins were observed in the elutions with NifQ (Fig. 4A, B). In these interactions, 85% of the total protein in the elution fractions corresponded to NifQ, and in the flowthrough an even larger amount to NifU was observed. Taken these proportions into consideration and measuring the iron content in the flowthrough and elution fractions, the iron:protein ratios could be determined. As shown in Figure 4C, incubation with NifU_S_ significantly increased the iron content in NifQ_H_ to around 2:1 molar ration when partnered with R-NifU_S_, and 1.5:1 with AS-NifU_S_. Similar results were observed when the interaction was carried out for 120 min, which led to close-to-saturation iron levels in NifQ_H_ when combined with R-NifU_S_ (Fig. S3).

**Figure 4.**
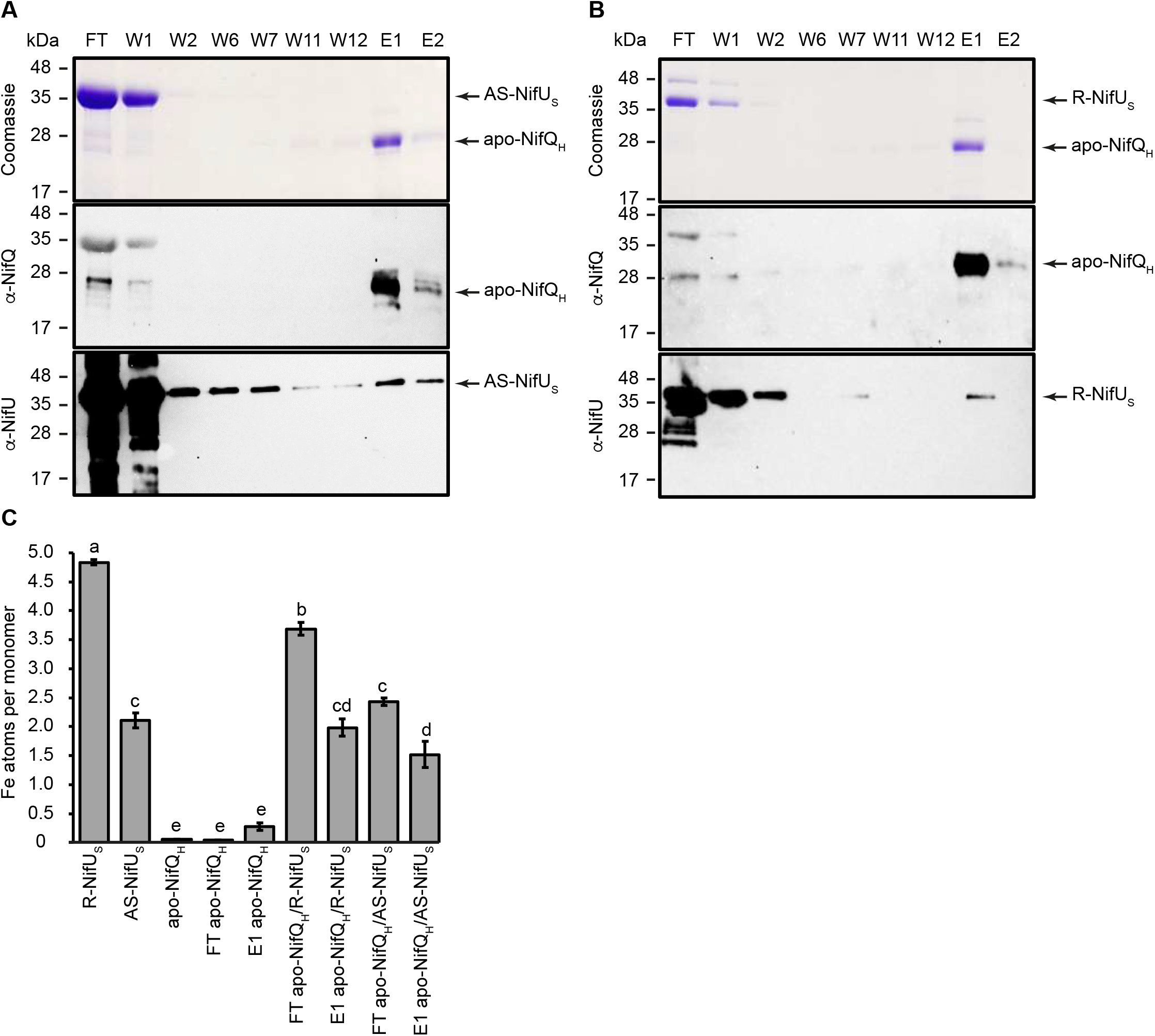
NifU_S_ transfers iron to apo-NifQ_H_. **A**. Top panel shows the Coomassie staining of an SDS-PAGE of flowthrough (FT), wash (W1-W12) and elution (E1-E2) fractions of a mixture solution containing AS-NifU_S_ and apo-NifQ_H_ passed through a Ni^2+^ column. The remaining panels are the immunoblots of the same fractions developed with anti-NifQ or an anti-NifU antibodies. **B**. Top panel shows the Coomassie staining of an SDS-PAGE of flowthrough (FT), wash (W1-W12) and elution (E1-E2) fractions of a mixture solution containing R-NifU_S_, and apo-NifQ_H_ passed through a Ni^2+^ column. The remaining panels are the immunoblots of the same fractions developed with anti-NifQ or an anti-NifU antibodies. Uncropped immunoblots and gels are shown in Supplementary Fig. 7. **C**. Iron content per monomer of pure isolated proteins, and in the FT and E1 fractions obtained from a passing through a Ni^2+^-column a solution in which apo-NifQ_H_ was incubated for 5 min with either AS-NifU_S_ or R-NifU_S_. Bars represent the average ± SD (n=2). Different letters indicate statistically significant differences (p < 0.01).

Iron binding to apo-NifQ could be due to sequestering the iron that may dissociate from NifU, instead of being the consequence of direct protein-protein transfer. If this were the case, separating the two proteins with a membrane that only allowed for iron diffusion but prevented the passage of the proteins, should still result in iron binding to apo-NifQ. However, when this control was carried out, using R-NifU_S_ or AS-NifU_S_, no iron was detected in the compartment containing apo-NifQ_H_ even when 120 min was allowed for iron to dissociate and diffuse (Fig. 5).

**Figure 5.**
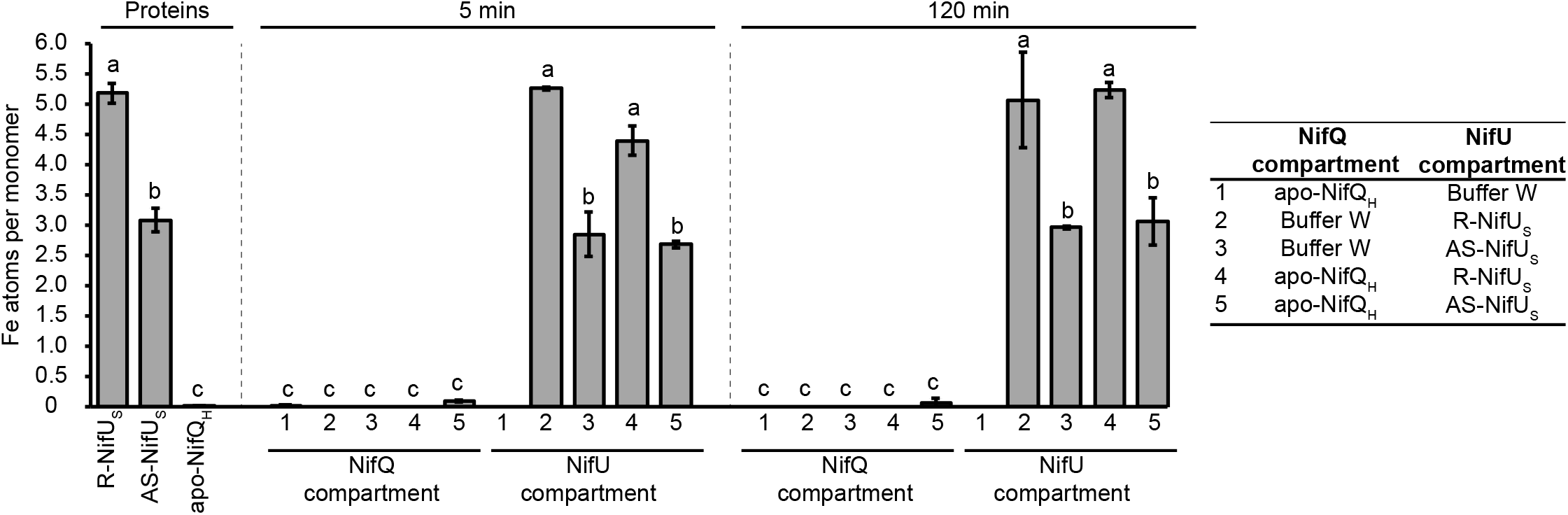
NifQ requires physical interaction from NifU to receive iron. Iron content per monomer of pure isolated proteins or from proteins separated by a 2-kDa pore-size cutoff dialysis membrane after 5 or 120 min incubation. Bars represent the average ± SD (n=2). Different letters indicate statistically significant differences (p < 0.01).

### Changes in the UV-visible absorption spectrum of NifQ upon interaction with NifU

The presence of [Fe-S] clusters in a protein affects its UV-visible signature. Both R-NifU_S_ and AS-NifU_S_ presented UV-vis absorption spectra characteristic of carrying O_2_-sensitive [Fe-S] cluster, with a peak around 330 nm and another around 420 nm (Fig. 6A). These peaks were not observed in apo-NifQ_H_ UV-vis absorption spectra, indicating the absence of any [Fe-S] cluster. Fractions obtained after the apo-NifQ_H_/R-NifU_S_ interaction were analyzed to determine their UV-vis absorption spectra. As shown in Figure 6B, the absorption spectra from the elution fraction, where more than 85% of the total proteins corresponded to NifQ_H_, presented the typical shoulder around 400 nm related to [Fe_4_-S_4_] cluster (8).

**Figure 6.**
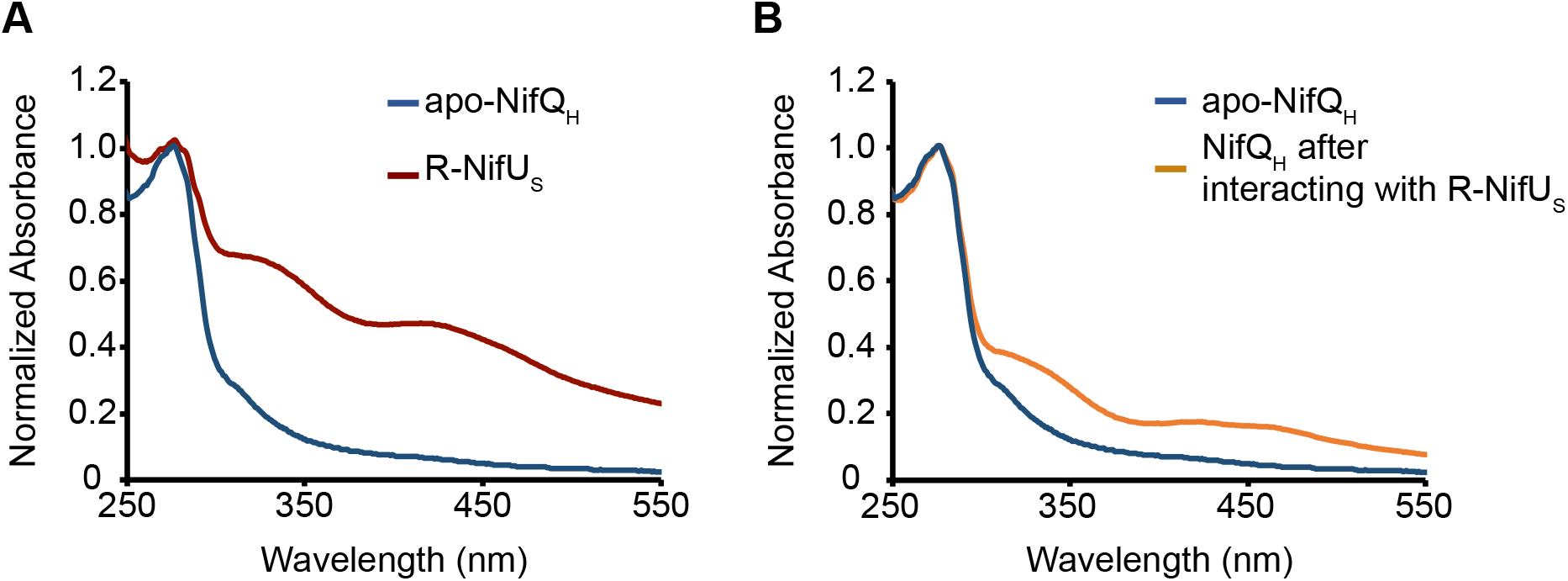
NifQ receives a [Fe_4_-S_4_] cluster from NifU. **A**. UV-vis spectra of pure proteins. **B**. UV-vis spectra of E1 elution after the apo-NifQ_H_/R-NifU_S_ interaction for 5 min. The absorbances were normalized at 280 value (*n*=3).

## Discussion

[Fe-S] proteins are present in all three domains of life, participating in a wide array of physiological processes that include DNA metabolism, energy transduction, or metabolic pathways (17). It has been estimated that 1-5 % of bacterial proteins contain some [Fe-S] cluster (18). All these groups can be made *in vitro* by simply providing iron and sulfur, while *in vivo* they are synthesized over protein scaffolds in a process requiring multiple enzymes (17). The importance of these scaffold proteins is evidenced by their essential nature for cell metabolism and their specialization for different metabolic processes. Furthermore, to date there is no known *de novo* synthesis of [Fe-S] groups directly in target apo-proteins. Consequently, a “bucket-brigade” of proteins direct the newly produced [Fe-S] clusters to the acceptor proteins (5, 17, 19). In this context, it is worth noting that the model diazotroph *A. vinelandii* uses NifU as the primary scaffold for [Fe-S] synthesis and transfer to nitrogenase components (7, 10).

Molybdate use for FeMo-co synthesis *via* NifQ requires of a pre-existing [Fe_3_-S_4_]^+^ group (14, 16). NifU and NifQ interaction when heterologously produced in *E. coli*, or when they are purified and combined under *in vitro*, conditions suggest that NifU could be the source of the [Fe_4_-S_4_] precursor. This is further supported by the increased iron content in NifQ when incubated with NifU, as well as by the spectroscopic signature of the repurified NifU-interacting NifQ which presents the absorbance pattern of [Fe_4_-S_4_] groups, characterized by a pronounced shoulder around 400 nm (8). The [Fe_4_-S_4_] cluster would then have to lose one iron, to create the [Fe_3_-S_4_]^+^ group required for molybdate to form the [Mo-Fe_3_-S_4_] prior to molybdenum transfer to the NifEN/NifH complex (14). It is to be expected that additional proteins will establish complexes with NifQ to mediate molybdate transfer and inclusion into the cluster.

Functional interaction between two proteins can sometimes evolve into one single protein with two different domains, each one of them corresponding to one of the original enzymes. This is advantageous to channel the product of one enzyme to the next, reducing diffusion times, increasing local substrate concentrations, and improving kinetics overall (20). *A. vinelandii* NifU is an example of this domain evolution since it combines an N-terminal IscU scaffold motive, a central ferredoxin fold, and a C-terminal NfuA-like domain (7). Another protein essential for nitrogenase maturation, NifB can also be found as standalone radical *S*-adenosylmethionine (SAM) domain that then interacts with the NifB-cofactor carrier NifX, or as combination of both proteins in a single polypeptide (21). Consistent with an adaptation to optimize protein-protein interactions, some delta-proteobacteria (such as *Geoalkalibacter ferrihydricus* or *Malonomonas rubra*) contain NifQ as the C-terminal domain of a larger protein that also includes an N-terminal domain with high homology to IscU/NifU proteins. Interestingly, the additional domain only shares homology with the N-terminal region of *A. vinelandii* NifU, what could indicate that the [Fe_4_-S_4_] clusters transferred to NifQ would mainly be synthesized in N-terminal NifU.

The interaction between two proteins exchanging substrates must be a relatively fast and labile process for it to work at optimal conditions and limit the subset of proteins and substrates lost in unproductive interactions. For instance, Cu^+^-chaperone CopZ rapidly dissociates from Cu^+^-transporting ATPase CopA after transferring Cu^+^ to prevent the apo-chaperone blocking the transfer site (22, 23). NifQ-NifU interaction is similarly conditional to the metalation state of NifQ. When NifQ already contains an [Fe-S] cluster -the holo-NifQ used in this study-the interaction with NifU does not occur or is severely weakened.

Beyond the specific metalation of NifQ, its interaction with NifU and the likely dependency on NifU activity signals the position in which iron and molybdenum metabolism for biological nitrogen fixation are coordinated. Only when sufficient iron is allocated for NifU, molybdenum might be used for NifQ. This co-regulation of both elements is also present in other molybdenum-dependent reactions, such as the synthesis of the molybdenum-cofactor (24).

In summary, these data indicate that NifU transfers [Fe_4_-S_4_] to all three major sets of [Fe-S] proteins in biological nitrogen fixation: NifH, NifB, and now also NifQ. This information is relevant as nitrogenase elements are being introduced and expressed in plants towards developing nitrogen-fixing crops. Co-expression with NifU has already shown to be essential for NifH activity when purified from plant chloroplasts (25), as well as for NifB obtained from yeast mitochondria (26). Therefore, it should be expected that similar co-expression with NifU would be needed for a functional NifQ-mediated molybdenum delivery pathway to nitrogenase in plants.

### Experimental Procedures

#### Escherichia coli strains and plasmids

*E. coli* strain BL21 (DE3) was used to express the proteins used in this study. The plasmid pN2LP30 was used to produce untagged NifU and NifS in *E. coli*. This plasmid was obtained by amplifying the *A. vinelandii nifUS* genes with primers 2495 and 2496 (Table S1) and using ELIC cloning (27) to introduce them in the *Nco*I/*Not*I digested pRSF*iscmetK*Duet-1 plasmid. To generate a NifQ_H_ expressing vector, the primers 1184 and 1185 (Table S1) were used to amplify the *nifQ* sequence from the *A. vinelandii* genomic DNA. The resulting amplicon was digested with *Pst*I and *Not*I and cloned into previously *Pst*I/*Not*I-digested pTrc99A. To produce _S_NifQ, the amplicon obtained from *A. vinelandii* genomic DNA using the primers NifQ-5’ and NifQ-3’ (Table S1) was digested with *Nde*I and *BamH*I and cloned in a similarly digested pT7-7 vector. Strep-tag was added to this vector by ligating at the *Nde*I site the overlapping oligonucleotides *Nde*I-Strep-tag-5’ and *Nde*I-Strep-tag-3’ (Table S1). The same procedure was used to fuse the Strep-tag to *nifS*-expressing vector pDB21223 (9). _H_NifU was obtained from cells transformed with plasmid pRHB609 (28). To produce NifU_S_, a 174 DNA fragment containing the last 99 nucleotides of *NifU* fused to the Strep-tag was synthesized (Integrated DNA Technologies, Coralville, IA), digested with *Sac*I and *BamH*I, and ligated in similarly digested *NifU*-encoding plasmid pDB525 (7).

### Culture conditions for Nif protein expression in *E. coli*

In general, gene expression was induced with 1 mM isopropyl β-D-thiogalactoside (IPTG) in cells growing in LB media supplemented with 100 μg/ml ampicillin at OD_600_ ≈ 0.6. After 3 h of induction at 37ºC, cells were collected by centrifugation at 4,000 x *g* for 10 minutes. Cells producing NifU_S_ were grown in LB media supplemented with 100 μg/ml ampicillin, with 0.2 mM ferric ammonium citrate and 2 mM L-cysteine. Induction was done at OD_600_ ≈ 0.6 with 0.5 mM IPTG for 5-6 hours at 37ºC. _H_NifU induction was performed at OD_600_ ≈ 0.7 with 1 mM IPTG and 0.1 mg/l Fe(NH_4_)_2_(SO_4_)_2_ for 14 hours at 18 ºC and 150 rpm.

#### Protein purification

Strep-tagged proteins were purified by Strep-Tactin XT affinity chromatography (SATC). Approximately 15-20 g of recombinant *E. coli* BL21(DE3) cells were resuspended for 30 minutes in 80 ml of lysis buffer A containing 50 mM Tris-HCl pH 8.0, 100 mM NaCl and 10% glycerol, 1 mM phenylmethylsulfonyl fluoride (PMSF). Cells were lysed in a French Press cell at 1,500 lb per square inch. The cell-free extract (CFE) was obtained after removing cell debris by centrifugation at 63,000 *x g* for 1 hour at 4°C and filtration with 0.45 μm pore size syringe filters (Sartorius). CFE was loaded onto a 1 ml Gravity flow Streptactin-XT high-capacity column (IBA Lifesciences), previously equilibrated with buffer A. The column was then washed 5 times with 2 column volumes (CV) of buffer A per wash. Bound protein was eluted in three steps with 1, 4 and 2 CV of buffer A containing 50 mM biotin per step.

His-tagged proteins were purified by Ni-NTA affinity chromatography. Approximately 20-25 g of recombinant *E. coli* BL21(DE3) cells were resuspended for 30 minutes in 100 ml of lysis buffer W 100 mM Tris-HCl pH 8.0, 150 mM NaCl and 10% glycerol, 1 mM PMSF. Cells were lysed and CFE was obtained as described above. CFE was loaded onto a 2-ml Ni-NTA Agarose column (Qiagen) equilibrated with buffer W supplemented with 5 mM imidazole. Column was washed 6 times with 1 CV of buffer W with 5 mM imidazole and 6 times with 1 CV of buffer W with 20 mM imidazole per wash. Protein was eluted buffer W containing 150- and 300-mM imidazole.

Apo-NifQ purifications that were carried out in aerobic conditions to promote losing any bound iron. All other proteins were purified under anaerobic conditions (< 5.0 ppm O_2_) inside a glovebox (COY Laboratories) using buffers previously made anaerobic by sparging with N_2_ overnight. Purification fractions were analyzed by electrophoresis.

Elution fractions were concentrated with 10-kDa cut-off pore size centrifugal membrane devices (Amicon Ultra-15, Millipore). Centrifugation procedure was performed at 4,000 x *g* for 45 min and this step was repeated until estimated biotin or imidazole concentration was lower than 50 nM and 500 nM, respectively. Protein concentration was determined by the bicinchoninic acid method (Pierce) with bovine serum albumin as the standard (29). For iron determination, the rapid colorimetric micro-method for the quantitation of complexed iron in biological samples was performed (30). Purified proteins were frozen and stored in liquid N_2_.

### In vitro *[Fe-S] cluster reconstitution of NifU*

Strep-tagged NifU purified from *E. coli* was reconstituted *in vitro* as described (31) with slight modifications. 20 μM of NifU dimer was prepared in 100 mM MOPS (pH 7.5) buffer containing 8 mM 1,4-dithiothreitol (DTT) and incubated at 37 °C for 30 min. To this mixture, 1 mM L-cysteine, 1 mM DTT, 225 nM NifS and 0.3 mM (NH_4_)_2_Fe(SO_4_)_2_ were added. Iron additions were divided in three steps of 15 min each until reaching the final concentration of 0.3 mM. The reconstitution mixture was kept in ice for 3 h and then desalted using 10-kDa cutoff pore size centrifugal membrane devices (Amicon, Millipore) to remove excess reagents. R-NifU protein was stored in liquid nitrogen until use.

#### Protein-protein interaction assays

Interaction assays were carried out for 5 min unless otherwise stated using 10 nmol of each protein in a glovebox (COY Laboratories) under anaerobic conditions. Those involving apo-_S_NifQ took place in Buffer A. _S_NifQ and its interacting proteins were recovered passing the solution through a 200 μl Gravity flow Streptactin-XT column (IBA Lifesciences), previously equilibrated with anaerobic buffer A. Column was washed 5 times with 2 CV of buffer A. The elution of target proteins from the resin was carried out by applying 0.5 CV, 1.4 CV and 0.8 CV of 2.5 mM desthiobiotin in Buffer A. When using NifQ_H_ as bait, the interaction was carried out in Buffer W. Proteins were separated using a 200 μl Ni-NTA agarose column equilibrated with anaerobic 5 mM imidazole in buffer W. The column was washed 6 times with 2 CV of 5 mM imidazole in buffer W and 6 times with 2 CV of 20 mM imidazole in buffer W per wash. Elution was performed with 150 mM imidazole in buffer W.

To assess the interaction between holo-_S_NifQ and AS-_H_NifU, 10 nmol of holo-_S_NifQ were immobilized on a 200 μl Gravity flow Streptactin-XT column, previously equilibrated with anaerobic buffer A. Column was washed twice with 2 CV of buffer A and 10 nmol of AS-_H_NifU were loaded onto the holo-_S_NifQ-charged column. This column was washed 3 times with 3 CV of buffer and eluted with 50 mM biotin in buffer A.

To test the diffusion of iron, 50 nmol of apo-NifQ_H_ and 50 nmol of R-NifU_S_ were incubated for 5 and 120 min inside an anaerobic glovebox (COY Laboratories), separated by inserting a 2-kDa pore-size cutoff dialysis membrane, previously equilibrated for 1 h with buffer W. Controls with only apo-NifQ_H_ on R-NifU_S_, AS-NifU_S_ on the other side of the membrane were carried out at the same time. At the indicated times, samples from both membrane sides were collected to determine the protein and iron concentration.

Protein content in all selected fractions was analyzed by SDS-PAGE using 12 % acrylamide/bisacrylamide (37.5:1) gels and visualized by Coomassie Brilliant Blue staining (32). For immunoblot analysis, proteins were transferred to nitrocellulose membranes for 45 min at 20 V using a Transfer-Blot® Semi Dry system (Bio-Rad). Immunoblot analyses were carried out with antibodies raised against *A*.*vinelandii* NifQ (1:2,500 dilution), NifU (1:2,500 dilution) and NifS (1:1,500 dilution) (29). A horseradish peroxidase conjugated anti-rabbit antibody (Invitrogen) diluted 1:15,000 was used as a secondary antibody. Chemiluminescent detection was carried out according to Pierce ECL Western Blotting Substrate kit’s instructions (ThermoFisher Scientific) and developed in an iBright FL1000 Imaging System (ThermoFisher Scientific).

#### Ultraviolet-visible spectroscopy

UV-visible absorption spectra were collected under anaerobic conditions (< 0.1 ppm O_2_) inside a glovebox (MBraun) in septum sealed-cuvettes to avoid the O_2_ contamination during the measurements in the Shimadzu UV-2600 spectrophotometer. Absorption (225 nm to 800 nm) was recorded, and the data were normalized to absorption at 280 nm.

#### Statistical methods

SPSS software (Statistical Package for Social Sciences) was used for statistical analyses. The data were compared using one way analyses of variance (ANOVA) followed by Bonferroni’s multiple comparation test (p< 0.01).

## Data Availability

The authors declare that the data supporting the findings of this study are available within the article, its supplementary information and data, and upon request.

## Supporting Information

This article contains supporting information.

## Acknowledgements

The authors would like to acknowledge Dr. Isidro Abreu (CBGP, UPM-INIA/CSIC) for his help in the protein-protein interaction assays, Dr. Lucía Payá (CBGP, UPM-INIA/CSIC) for providing the PN2LP30 vector, and Dr. Dennis Dean and Ms. Valerie L. Cash (Virginia Tech) for their gift of the _S_NifQ, NifU_S_, and _S_NifS expressing plasmids.

## Funding and Additional Information

This work was supported in part by the Bill & Melinda Gates Foundation (INV-005889). Under the grant conditions of the Foundation, a Creative Commons Attribution 4.0 Generic License has already been assigned to the Author Accepted Manuscript version that might arise from this submission. EB was funded by the Severo Ochoa Programme for Centres of Excellence in R&D from Agencia Estatal de Investigación of Spain (grant SEV-2016-0672) received by Centro de Biotecnología y Genómica de Plantas (UPM-INIA/CSIC). XJ is recipient of a doctoral fellowship from Universidad Politécnica de Madrid.

## Author contribution

EB performed most of the experiments. XJ prepared the reconstituted NifU. EJV carried out the holo-NifQ purifications. EB, LMR and MGG designed experiments, analyzed data, and wrote the manuscript with input from the other authors.

## Figure Legends

**Figure S1. NifU and NifS do not bind to the Streptactin resin**. Top panel shows the Coomassie staining of an SDS-PAGE of cell free extract (CFE), flowthrough (FT), wash (W1-W6) and elution (E1-E3) fractions of *nifU* and *nifS*-expressing *E. coli* extracts passed through a Streptactin column. The remaining panels are the immunoblots of the same fractions developed with anti-NifU or an anti-NifS antibodies. Images show a representative assay (*n*=2). Uncropped immunoblots and gels are shown in Supplementary Fig. 8.

**Figure S2. AS-NifU**_**S**_ **and** _**S**_**NifS proteins do not bind to a Ni**^**2+**^ **column**. Top panel shows the Coomassie staining of an SDS-PAGE of flowthrough (FT), wash (W1-W12) and elution (E1-E2) fractions of a mixture solution containing AS-NifU_S_ and _S_NifS passed through a Ni^2+^ column. The remaining panels are the immunoblots of the same fractions developed with anti-NifU or an anti-NifS antibodies. Images show a representative assay (*n*=3). Uncropped immunoblots and gels are shown in Supplementary Fig. 9.

**Figure S3. NifU**_**S**_ **transfers iron to apo-NifQ**_**H**_. **A**. Top panel shows the Coomassie staining of an SDS-PAGE of flowthrough (FT), wash (W1-W12) and elution (E1-E2) fractions of a mixture solution containing AS-NifU_S_ and apo-NifQ_H_ passed through a Ni^2+^ column. The remaining panels are the immunoblots of the same fractions developed with anti-NifQ or an anti-NifU antibodies. **B**. Top panel shows the Coomassie staining of an SDS-PAGE of flowthrough (FT), wash (W1-W12) and elution (E1-E2) fractions of a solution mixture containing R-NifU_S_ and apo-NifQ_H_ passed through a Ni^2+^ column. The remaining panels are the immunoblots of the same fractions developed with anti-NifQ or an anti-NifU antibodies. Uncropped immunoblots and gels are shown in Supplementary Fig. 10. **C**. Iron content per monomer of pure isolated protein, and in the FT and E1 fractions obtained from a passing through a Ni^2+^-column a solution in which apo-NifQ_H_ was incubated for 120 min with either AS-NifU_S_ or R-NifU_S_. Bars represent the average ± SD (n=2). Different letters indicate statistically significant differences (p < 0.01).

**Figure S4**: Uncropped Coomassie-stained gels shown in Figure 1 (A). Uncropped immunoblots shown in Figure 1 that correspond to immunoblotting with an anti-NifQ antibody (B), an anti-NifU antibody (C) or an anti-NifS antibody (D).

**Figure S5**: Uncropped Coomassie-stained gels shown in Figure 2 (A). Uncropped immunoblots shown in Figure 2 that correspond to immunoblotting with an anti-NifQ (B), an anti-NifU (C), or an anti-NifS (D) antibodies.

**Figure S6**: Uncropped Coomassie-stained gels shown in Figure 3 (A). Uncropped immunoblots shown in Figure 3 that correspond immunoblotting with an anti-NifQ (B), or an anti-NifU (C) antibodies.

**Figure S7**: Uncropped Coomassie-stained gels shown in Figure 4A (A). Uncropped immunoblots shown in Figure 4A that correspond immunoblotting with an anti-NifQ (B), or an anti-NifU (C) antibodies. Uncropped Coomassie shown in Figure 4B (D). Uncropped immunoblots shown in Figure 4B that correspond immunoblotting with an anti-NifQ (E), or an anti-NifU (F) antibodies.

**Figure S8**: Uncropped Coomassie-stained gels shown in Figure S1 (A). Uncropped immunoblots shown in Figure S1 that correspond immunoblotting with an anti-NifU (B), or an anti-NifS (C) antibodies.

**Figure S9**: Uncropped Coomassie-stained gels shown in Figure S2 (A). Uncropped immunoblots shown in Figure S2 that correspond immunoblotting with an anti-NifU (B), or an anti-NifS (C) antibodies.

**Figure S10**: Uncropped Coomassie-stained gels shown in Figure S3A (A). Uncropped immunoblots shown in Figure S3A that correspond immunoblotting with an anti-NifQ (B), or an anti-NifU (C) antibodies. Uncropped Coomassie shown in Figure S3B (D). Uncropped immunoblots shown in Figure S3B that correspond immunoblotting with an anti-NifQ (E), or an anti-NifU (F) antibodies.

**Table S1**: Primers used in this study.

