## Supplementary figures and images for "*Azotobacter vinelandii* scaffold protein NifU transfers iron to NifQ as part of the iron-molybdenum cofactor biosynthesis pathway for nitrogenase"

### Supporting Figures

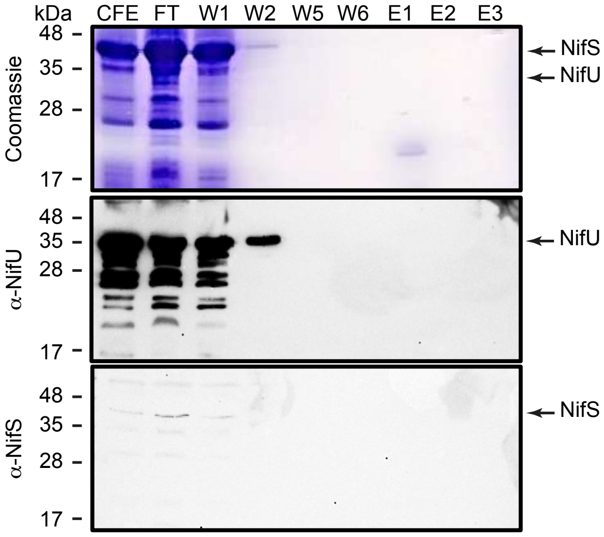

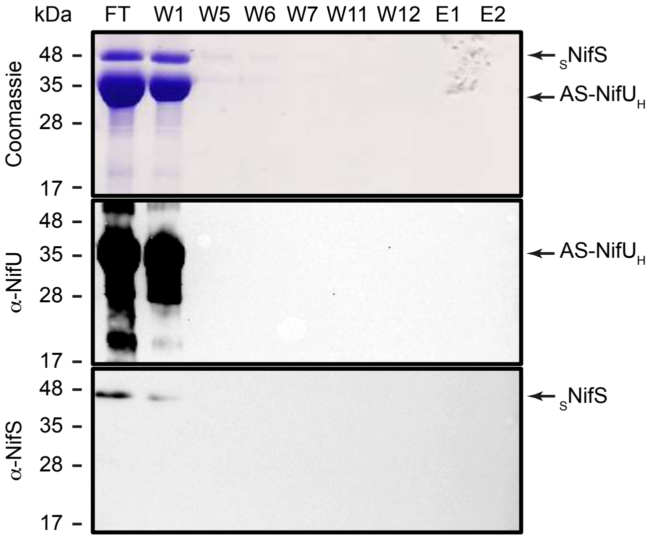

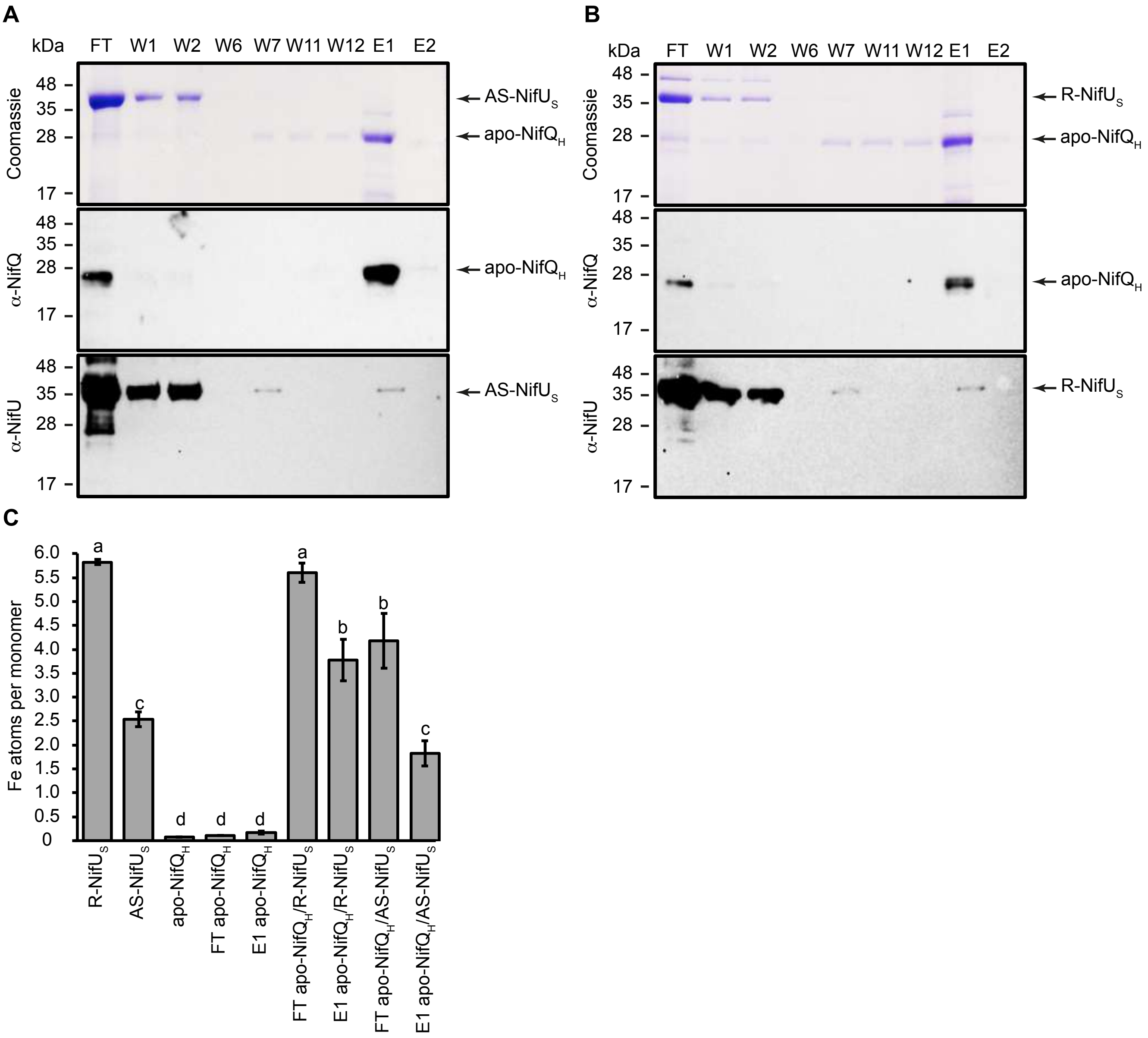

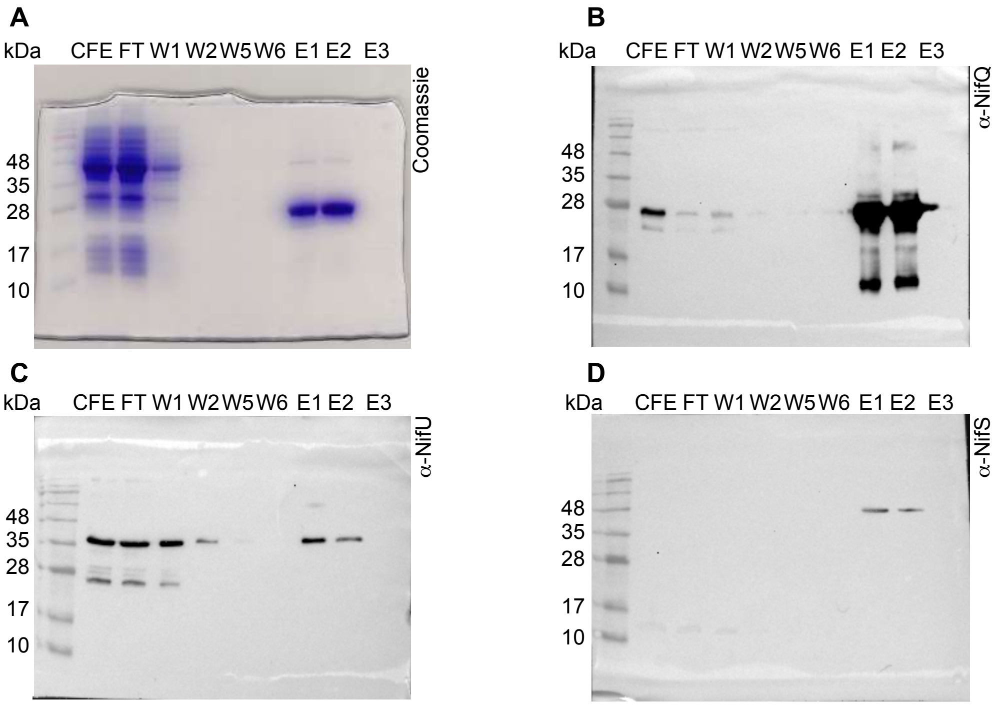

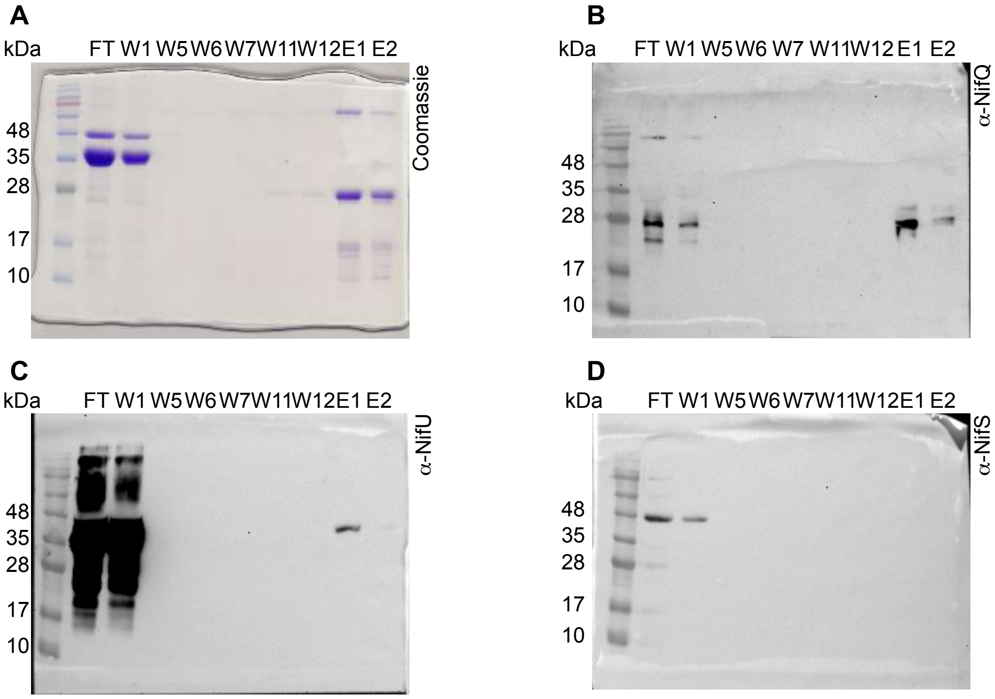

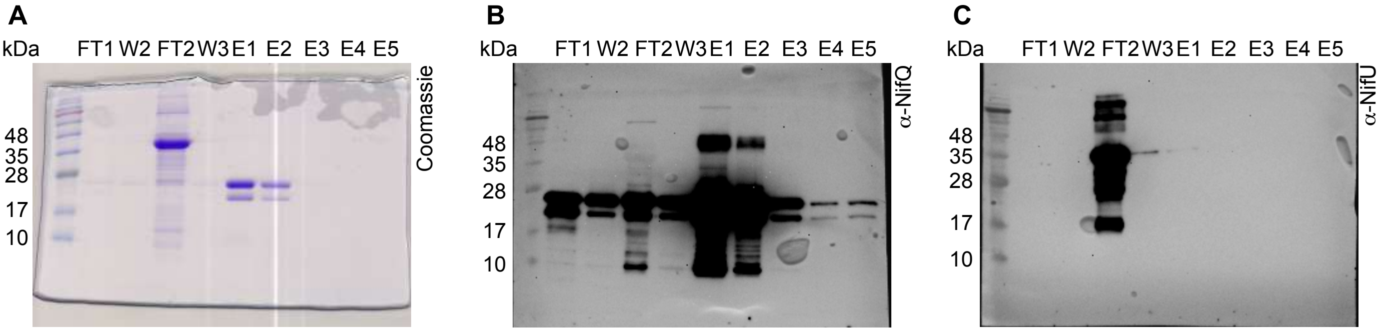

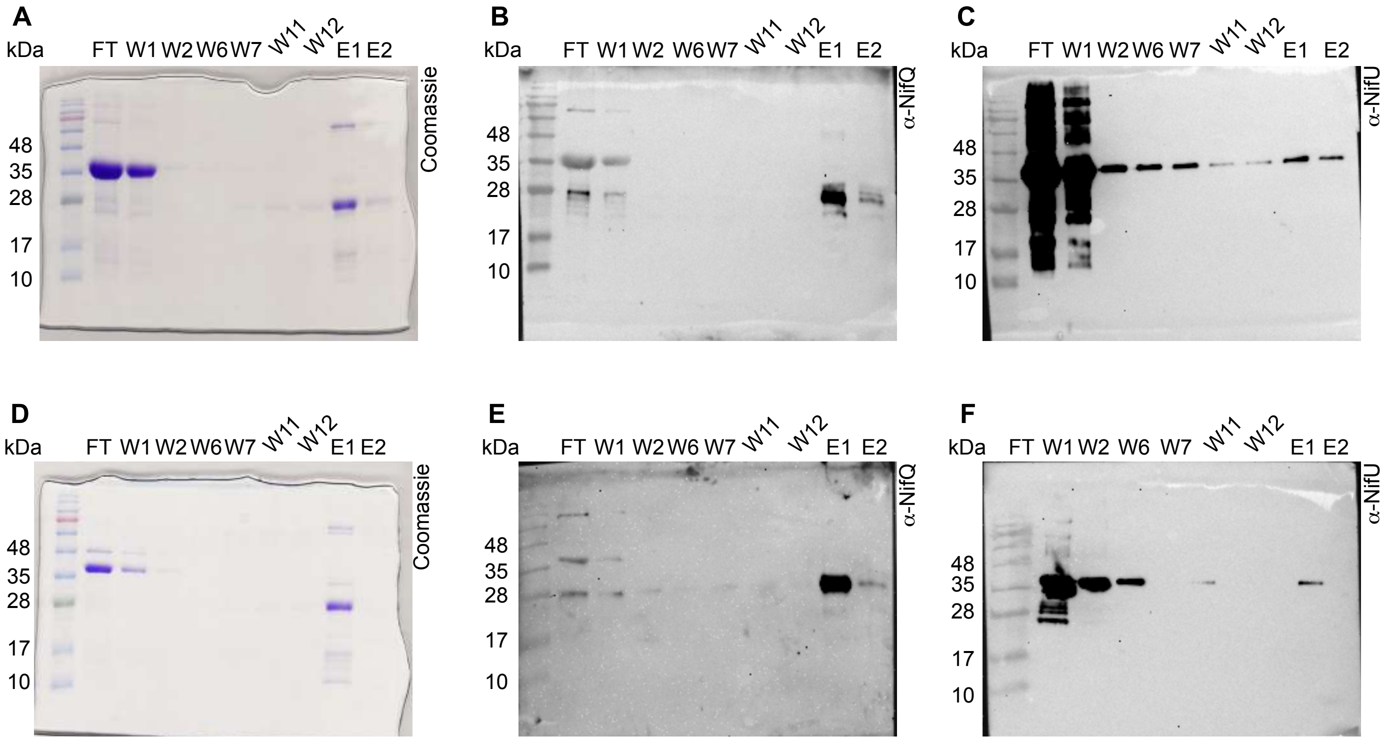

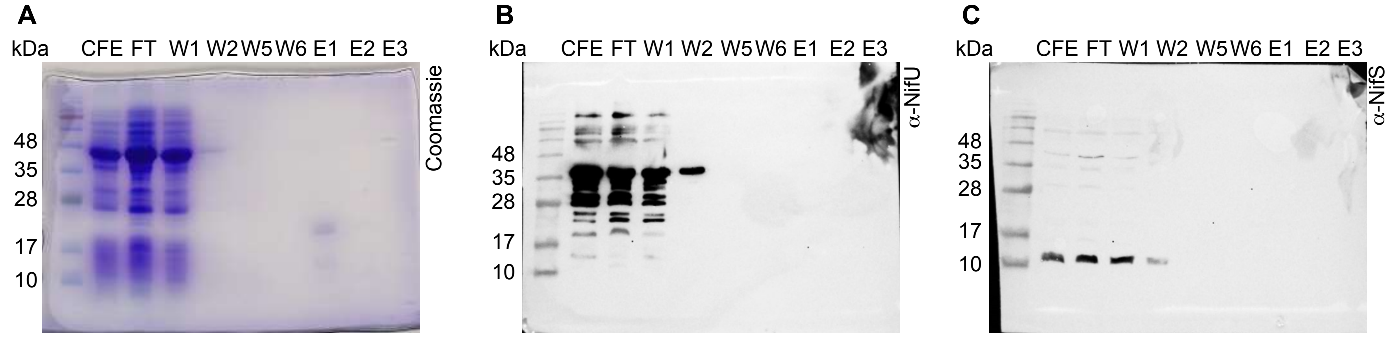

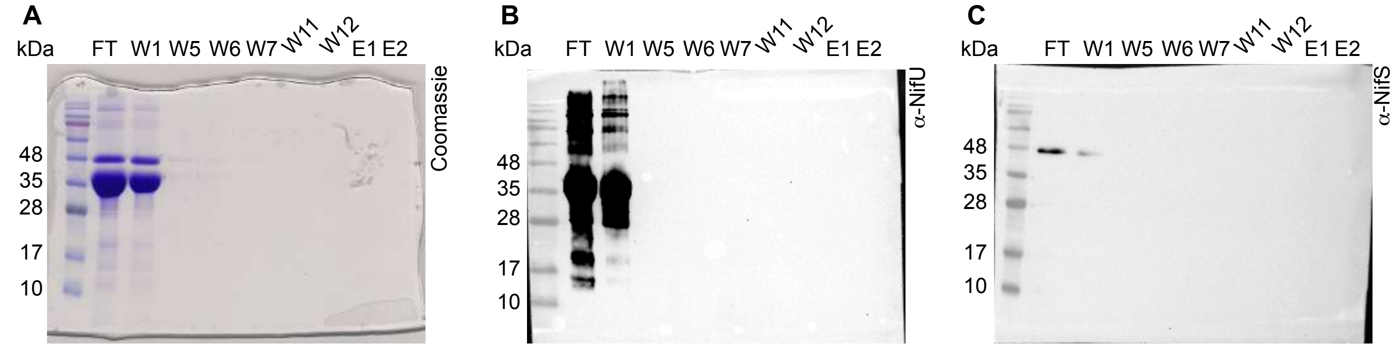

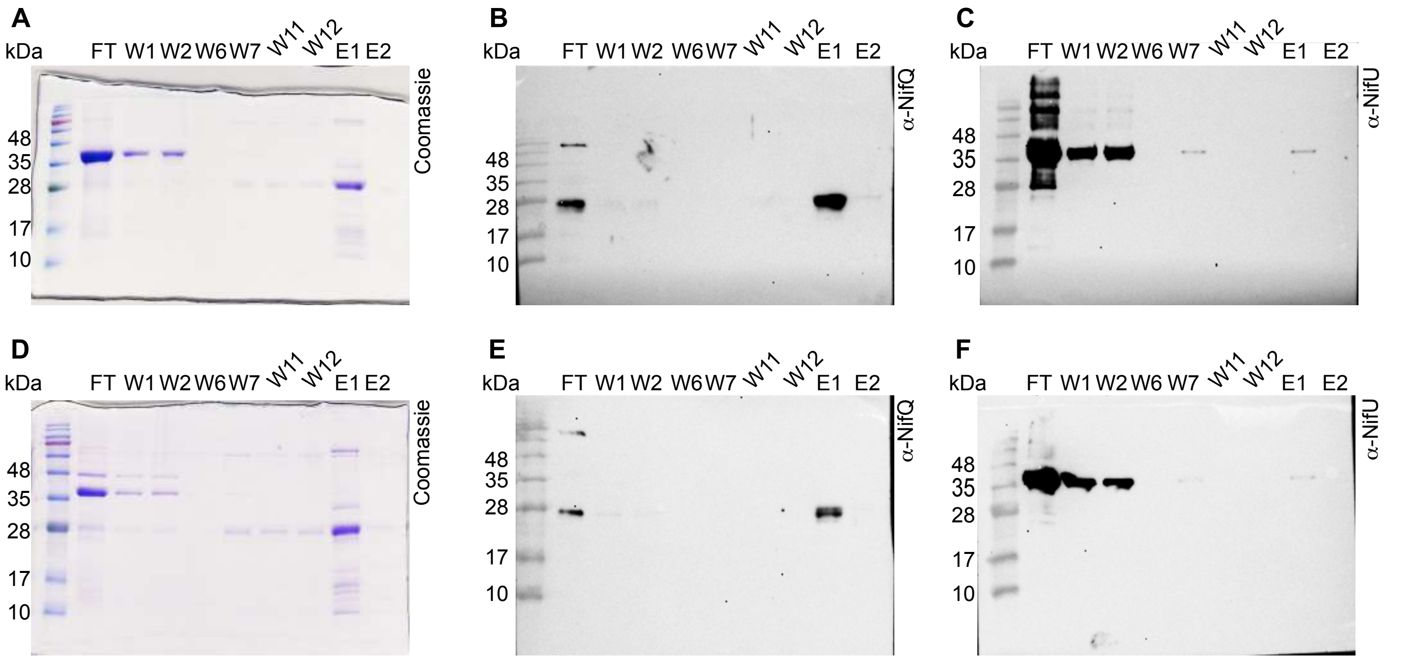
