## Supporting Table for "*Azotobacter vinelandii* scaffold protein NifU transfers iron to NifQ as part of the iron-molybdenum cofactor biosynthesis pathway for nitrogenase"

**Table S1.** Primers used in this study.

| Name | Sequence | Use |
|---|---|---|
| 2495 | TTAATAAGGAGATATACCATGGCCTGGGATTATTCGGAAA | Cloning of *nifUS* in pN2LP30 |
| 2496 | TTCGACTTAAGCATTATGCGGCCGCTCAGCCGTAGACCGG | Cloning of *nifUS* in pN2LP30 |
| 1184 | AAATTCTGCAGATGGGCAGCGCCGCG | Generation of $NifQ_H$ |
| 1185 | AATTGCGGCCGCGAGAATCGGGTCATATCTCTGCTCC | Generation of $NifQ_H$ |
| nifQ-5’ | CATG CAT ATG GGC AGC GCC GCG GCC | Amplification of NifQ |
| nifQ-3’ | CTAC GGA TCC TGG CCG GCC AGC AGG | Amplification of NifQ |
| *Nde*I-Strep-tag -5' | TATGGCTAGCTGGAGCCACCCGCAGTTCGAAAAACA | Addition of Strep-tag to *Nde*I sites |
| *Nde*I-Strep-tag -3' | TATGTTTTTCGAACTGCGGGTGGCTCCAGCTAGCCA | Addition of Strep-tag to *Nde*I sites |
